# Interfacing Live Cells with Surfaces: A Concurrent Control Technique for Quantifying Surface Ligand Activity

**DOI:** 10.1101/2021.04.27.441513

**Authors:** Michael C. Robitaille, Joseph A. Christodoulides, Patrick Calhoun, Jeff M. Byers, Marc P. Raphael

## Abstract

Surface ligand activity is a key design parameter for successfully interfacing surfaces with cells - whether in the context of *in vitro* investigations for understanding cellular signaling pathways or more applied applications in drug delivery and medical implants. Unlike other crucial surface parameters, such as stiffness and roughness, surface ligand activity currently lacks a standardized measurement approach that can be readily paired with live cell investigations. To fill this void, we have developed a concurrent control technique for characterizing *in vitro* ligand surface activity. Pairs of gold-coated glass chips were biofunctionalized with RGD ligand in a parallel workflow: one chip for *in vitro* applications and the other for surface plasmon resonance (SPR) based RGD activity characterization. Recombinant α_V_β_3_ integrins were injected over the SPR chip surface as mimics of the cellular membrane bound receptors and the resulting binding kinetics parameterized to quantify ligand activity. These activity measurements were correlated with cell morphological features, measured by interfacing MDA-MB-231 cells with the *in vitro* chip surfaces on the live cell microscope. We show that the SPR concurrent control approach has multiple advantages based on the facts that SPR is a standardized technique and has the sensitivity to measure ligand activity across the most relevant range of extracellular surface densities. Furthermore, by pairing both SPR and *in vitro* approaches, a comparison of the results can provide biological insights into the nature of cellular adhesion and dynamics.

## Introduction

The disciplines of lithography and surface chemistry are now regularly combined to engineer surfaces capable of directing cellular decisions^1^. The cell line is chosen based on biological context: cancer cells for understanding metastasis-related migrations^2^, epithelial for the study of wound healing^3^, osteoblasts for adhesion to medical implants^4^, and so forth. The surfaces are designed to explore the causal effects of one or more engineered inputs, which can be either chemical, topographical or mechanical in nature. Four external inputs to which a wide range of cell lines have been shown to respond dynamically and morphologically with significant effect sizes are: (1) active surface ligand density^5-9^ (2) substrate stiffness^10- 13^ (3) surface topography^14-18^ and (4) ligand tether length^19,20^.

To vary one input parameter while holding the others constant is not sufficient for understanding cellular response unless the values of those constants are measured as well^21-23^. For instance, the statement that focal adhesion (FA) length has been shown to be independent of surface ligand density is only true in the context of stiff substrates such as glass. The FA length varies strongly on softer substrates, emphasizing that parameters such as stiffness require quantitative characterization as well as the variable ligand density^24^. In a similar manner, if stem cell differentiation is viewed in the context of substrate stiffness without measuring ligand density or roughness, an incomplete view emerges given that all these input parameters have been shown to direct differentiation to varying degrees^25,26^. Traditionally, experiments have not been conducted with a quantitative knowledge of where the cell-interfacing surface lies in this multi-dimensional parameter space, thus making it challenging to place the results in context and compare studies.

It is now understood that the cell interfacing surface must be viewed as an instrument capable of multi-cue cellular inputs, and as such, requires thorough calibration. Fortunately, most relevant surface parameters can be quantified using a range of surface metrology techniques, which we will refer to here as concurrent controls (also sometimes called witness samples) because they are conducted on surfaces fabricated in parallel with those used in the cell experiments but dedicated to surface characterization. The most broadly applicable of these techniques for surface characterization is atomic force microscopy (AFM). When used as a scanning probe to map the surface, the resulting surface topography data enables surface roughness quantification. Alternatively, if the AFM tip is pressed into a soft material (*e*.*g*. polydimethylsiloxane (PDMS), hydrogel) then the deflection curves, often fitted with the Hertz model, can be used to infer material stiffness. Finally, a biofunctionalized AFM tip can be slowly retracted from the surface after allowing for receptor-ligand binding to take place. In the case of single molecule binding events, the strongly nonlinear force curve reveals when the ligand tether has reached its maximum extension. If AFM is not an option, there are numerous other techniques which work at larger length scales and give equally valuable insight into these surface properties: ellipsometry for surface roughness; rheometry for material stiffness; and surface force apparatus measurements for tether length.

Notably, there are no well-established techniques for independently characterizing surface ligand activity. This is despite the fact that determining surface-bound ligand activity is as much of interest as roughness, tether length and stiffness, if not more, since cell morphology and viability depend on it strongly. The challenge has been that the most commonly used techniques, such as the radio or fluorescence labeling of ligands, are proportional to ligand *density* rather than ligand *activity*, and the two values can be strikingly different. For common assays like ELISA, for example, it is estimated that over 90% of surface conjugated antibodies are inactive for a variety of reasons, including blocked binding motifs, partial denaturation and inaccessible orientations of the antigen binding site^27-29^.

An important technological advance toward overcoming this roadblock has been to physically define the surface density by using gold nanoparticles or lithographically patterned gold nanostructures with well-defined spacings^6,9,24^. When conjugated with small peptide ligands such as RGD, which are less prone to denaturation and more readily orientated, the result is well quantified ligand densities with increased likelihood of actively engaging the cell receptor. We emphasize the word *active* in this work because we have previously shown that RGD surface activity can be dramatically reduced by blocking from a variety of common media and surface functionalization components (*e*.*g*. serum, BSA and PLL-PEG), which in turn affects cell phenotype^30^. Thus, even when using sophisticated lithographic approaches, the <u>active</u> ligand density can vary with laboratory methodology. Furthermore, these lithographic techniques are not commercially available, and their fabrication involves skill sets not common to typical cell biology laboratories.

Here we introduce a concurrent control technique for characterizing *in vitro* ligand surface activity utilizing surface plasmon resonance (SPR) on planar substrates. A substrate design consisting of glass coated with thin film gold (Au) is used for both the live cell microscopy chips and the SPR chips, enabling identical surface functionalization protocols to be applied to both. For the SPR chip, recombinant receptor proteins, specifically recombinant α_V_β_3_ integrins, are injected over the surface as mimics of the cellular membrane bound receptors and the resulting binding kinetics parameterized to characterize the ligand activity. Live cell experiments are conducted on the in-parallel produced *in vitro* chip – an approach analogous to other *in vitro* surface characterization workflows such as those used for surface roughness or stiffness. The technique described here has multiple advantages: (1) receptor-ligand kinetics as characterized by SPR has been demonstrated to be reproducible across SPR platforms and laboratories, thereby enabling the standardization of surface activity measurements^31^; (2) we show the SPR technique has the sensitivity to measure ligand activity across the range of surface densities necessary to trigger cell adhesion and commonly encountered in the extracellular environment; (3) the SPR chip offers a straightforward means of not only characterizing ligand activity but also whether culture media or other added factors degrade ligand activity, and if so, by how much^30^; (4) we demonstrate that when ligand spacing studies are investigated using both SPR and *in vitro* approaches, a comparison of the results can provide biological insights into the nature of cellular adhesion and dynamics.

## Materials & Methods

Recombinant human α_V_β_3_ integrins (R&D Systems, #3050-AV-050) were reconstituted in 20 mM Tris buffered saline (TBS), pH 7.4, and 0.9% NaCl (Sigma, #T5912). The RGD-based peptide cRGDfK (cRGD, Peptides International, #PCI-3661-PI) was reconstituted in modified Dulbecco’s phosphate buffered saline (DPBS, Thermo, #28344) titrated with sodium hydroxide to pH 8.0. The self-assembled monolayer (SAM) thiols, SH-(CH_2_)_11_-EG_5_-OH (SPO) and SH-(CH_2_)_11_-EG_6_-COH_2_-COOH (SPC) (Prochimia Surfaces Sp.) were reconstituted in 200 proof, anhydrous ethanol. N-hydroxysulfosuccinimide (sulfo-NHS) and 1-ethyl-3-[3-dimethylaminopropyl]carbodiimide hydrochloride (EDC) from Thermo Scientific were dissolved in 18 MOhm DDW immediately before use. Ethanolamine (Sigma, #E905) was diluted in pH 7.4 DPBS to 0.1M. Ultrapure sodium dodecyl sulfate (SDS) solution (Invitrogen, #15553027) was diluted with 18 MOhm DDW unless otherwise specified.

For SPR measurements, commercially available bare gold sensor chips were purchased from Cytiva (SIA Kit Au). For *in vitro* studies, 20 nm Au thin films with sub-nm surface roughness were deposited on 25.4 mm diameter, No. 1.5 glass coverslips by e-beam evaporation as previously described^30^. The gold film was thin enough to allow for phase contrast microscopy techniques to be employed but thick enough for a continuous film to be formed on the substrate.

The surfaces and thiol design specifications defined the roughness, stiffness and tether length parameter space of the experiments: the surfaces are ultra smooth with an average surface roughness of 0.5 nm based on AFM characterization^30^; the substrate stiffness is maximal due to direct conjugation of the cRGD peptide to the underlying Au/glass substrate (> 50 GPa versus ∼ 1kPa -1MPa for most hydrogel constructs); the tether length, based on the constituent alkane and PEG linker lengths, is 3.8 nm which is categorized as short for PEG tether linker length studies that typically run from several nanometers to hundreds on nanometers^19,32^. As will be discussed in detail below, the surface activity was systematically varied by adjustments of the SPO:SPC ratio.

### Surface Plasmon Resonance (SPR) Chip Functionalization

For SPR studies using the Biacore 8K instrument, SIA bare gold sensor chips were cleaned down to the bare gold surface by a reactive ion etch (RIE) system utilizing 5% hydrogen, 95% argon gas mixture with the pressure regulated at 300 mT and the RF power at 40W, and then functionalized with a two component SAM consisting of SPO and SPC (Prochimia Surfaces Sp.) as previously described.^33^ The chips were immersed for 18 hrs in an 0.5 mM ethanolic-based thiol solution (room temperature) with a SPO to SPC ratios of 25:1, 250:1, 2500:1, 25K:1 and 250K:1 to vary the average ligand spacing. The chips were then rinsed with EtOH, dried under flowing nitrogen gas and mounted atop the Biacore SIA sensor inserts according to the manufacturer’s instructions. Activation of the SPC component with cRGD consisted of flowing a 33 mM: 133 mM ratio of sulfo-NHS:EDC for 5 min, followed by flowing 0.35 mg/mL cRGDfK for 5 min. Next, unreacted -COOH groups were blocked by flowing 0.1 M ethanolamine for 5 min. Finally, non-specifically bound reagents were removed by flowing 0.5% SDS (w/v, DDW) for 5 min. For all immobilization steps the flow rate was 10 µL/min and 18 MOhm DDW was used a running buffer. The Biacore 8K design incorporates 8 channels, each consisting of sensing lane paired with a reference lane for the subtraction of non-specifically bound analyte. The same immobilization protocol was used for both reference and sensing lanes with the exception that no cRGD was applied to the reference lane.

### SPR-Based Kinetic Rate Constants and Surface Activity Measurements

TBS + 0.5mM MnCl_2_ was used as running buffer for all kinetic rate constant and surface activity measurements with recombinant human α_V_β_3_ integrins introduced at a flow rate was 30 µL/min for all steps. Before the integrin introduction, the chip sensing surfaces were normalized using a 70% glycerol solution (BiaNorm, Cytiva) and then rinsed with 0.25% SDS (w/v, TBS) and multiple TBS rinses (400 sec each). All SPR experiments incorporated at least one control channel (blank) of only running buffer used to subtract out instrumental drift. The plotted SPR response curves have the reference lane and the buffer control channel subtracted.

### *In Vitro* Chip Functionalization and Cell Culture

As with the SPR chips, the Au coated coverslips were cleaned down to the bare gold surface using a RIE system with the same gas, pressure and power conditions described above and then functionalized with a two component self-assembled monolayer (SAM) by immersion for 18 h in an ethanolic-based thiol solution (0.5 mM). For the Au thin film biofunctionalization, the SPR self-assembled monolayer and immobilization procedures were repeated except all solutions were manually drop coated atop the chips and extensively washed with DDW. As a negative control for cell adhesion studies, additional *in vitro* chips were prepared consisting of only SPO thiols.

MDA-MB-231 cells (ATCC, #HTB-26) were cultured in DMEM supplemented with 10% fetal bovine serum (ATCC) and maintained at 37°C and 5% CO_2_. Prior to seeding on functionalized Au films, cells were harvested in their logarithmic growth phase (Supporting Information, Fig. S1) and trypsinized by adding 1 mL of trypsin-EDTA (ATCC) at 37°C for 3-4 min before the addition of DMEM (no serum). The cells were spun down at 125 × g for 5 min and the pellet re-suspended in 5 mL DMEM (no serum). Cells were spun down again and re-suspended in serum-free media to thoroughly eliminate all trypsin and serum components. As an additional negative control for the cell adhesion studies, the cells were resuspended in a DMEM-based solution containing 1 mM of cRGD for 60 min before drop coating on 25:1 thiol ratio chips. Cells exposed to cRGD in solution in this manner remained rounded and did not spread on the chips (data not shown).

### Live cell microscopy

Live cell imaging was performed using phase contrast illumination with a 10X, 0.9 numerical aperture objective. A heated stage and temperature controlled enclosure held the stage temperature at 37.0 ± 0.04°C (Zeiss). Humidity and CO_2_ were regulated at 98% and 5%, respectively. Cells were added to the functionalized chip to achieve cell densities of approximately 8 cells per field of view with image acquisition typically beginning within 20 min. Time-lapse, live-cell microscopy was performed every 10 minutes utilizing an automated X, Y stage to image 10-20 fields of view (FOV) per time point. Live cell imaging experiments were performed for at least 15 hours.

### Data Analysis

For every thiol ratio investigated, at least 3 SPR chips were independently prepared beginning with the cleaning of the bare Au surface, followed by ligand functionalization and integrin application. The multichannel results from the three chips were pooled to give at least five replicates for each thiol ratio.

For the *in vitro* studies, an experimental unit (*N* = 1) was defined as the FOV rather than a single cell due to occasional cell-to-cell interactions and mitosis events which prevented the cells from being counted as independent units. Each FOV in a given experiment was defined as a technical replicate and typically contained between five and ten cells. Morphological features (*e*.*g*. cell area, cell circularity) were first averaged within the FOV and then for all technical replicates. A biological replicate was defined as a completely new experiment from Au surface preparation to live cell microscopy. Three biological replicates were conducted per thiol ratio such that the pooling of technical replicate across biological replicates resulted in an *N* ≥ 30 for each thiol ratio.

To ensure objective live cell segmentation and analysis from the imagery, a completely automated self-supervised machine learning (SSML) algorithm was employed that required no curated training data or parameter tuning. For each image, the algorithm outputs metadata which includes the outlines of the segmented cells as well as morphological parameters such as cell area and circularity.^34^

## Results and Discussion

As with other *in vitro* concurrent control approaches (*e*.*g*. surface roughness, substrate stiffness), the goal was to prepare two sets of chips in parallel: one set for quantifying surface ligand activity and the other for cell biology applications. The left side of Fig. 1 shows an overview of the procedure in which the SPR and *in vitro* chip surfaces are simultaneously cleaned via plasma ashing and then simultaneously activated by the deposition of the two-component SAM. Because thin film Au surfaces are both live-cell compatible and standard use in commercial SPR instruments for biomolecular kinetic characterization, identical functionalization protocols can be applied using parallel processes. For the SPR chips, the cRGD conjugation took place in the Biacore 8K instrument in an automated fashion followed by the introduction of recombinant integrins for analysis; for the *in vitro* chips the identical cRGD conjugation protocol was carried out manually, followed by the addition of live cells and live cell microscopy time-lapse experiments.

**Figure 1.**
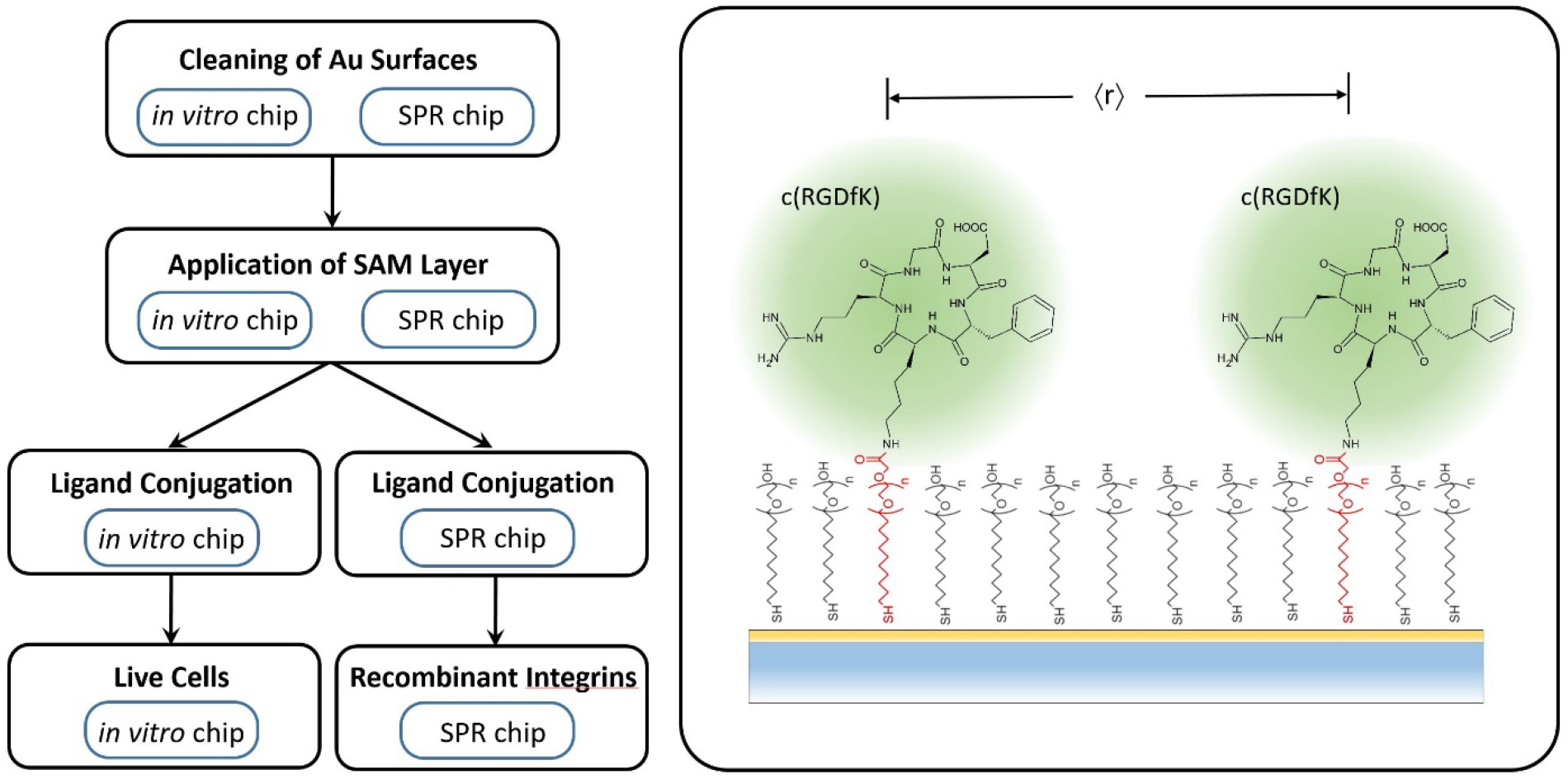
Workflow and surface chemistry. (left) The first two steps, cleaning and SAM layer activation, took place simultaneously for the *in vitro* and SPR chips. The next step, cRGD ligand conjugation, was conducted in an automated fashion in the Biacore 8K instrument for the SPR, while the identical protocol was carried out manually for the *in vitro* chips. The final step for *in vitro* chips was the adding of cells followed by time-lapsed live cell microscopy experiments; for the SPR chips the analogous step was the introduction of α_V_β_3_ recombinant integrins. (right) The spacing of cRGD ligands was controlled by varying the ratio of the hydroxyl-terminated spacer thiol molecules (SPO, black) to the carboxyl terminated thiol molecules (SPC, red) use for crosslinking the cRGD.

We first describe the procedure for characterizing the surface ligand activity using the concurrent control SPR chip. To quantify the cRGD-α_V_β_3_ kinetic rate constants, α_V_β_3_ concentrations ranging from 50 nM to 0.21 nM were microfluidically introduced over SPR sensor surfaces functionalized with cRGD (Fig. 2a). The one-to-one, first-order kinetic rate constant fits of *K*_*a*_ = 2.05 × 10^5^ ± 0.62 × 10^5^ *M*^−1^*s*^−1^ and *K*_*d*_= 4.60 × 10^−5^ ± 0.61 × 10^−6^ *s*^−1^ (*K*_*D*_ = 0.32 ± 0.08 *nM*) were consistent amongst 6 independently prepared chips across the range of thiol ratios. These rate constants serve as a receptor-ligand binding calibration, ensuring that surface activity measurements can be reliably compared from run to run and instrument to instrument.

**Figure 2.**
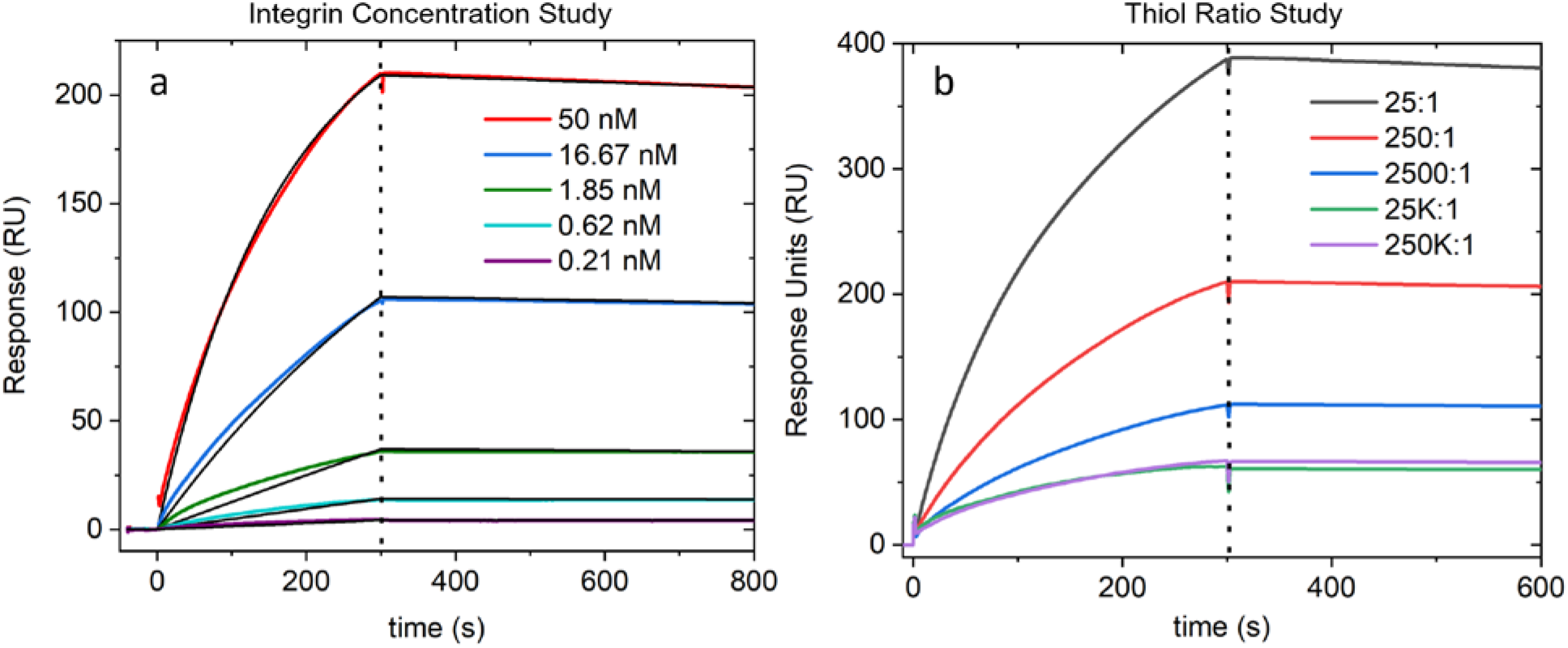
Recombinant α_V_β_3_ integrin/cRGD kinetics and instrument sensitivity studies. (a) Integrin concentration study (250:1 thiol ratio chip) in which [α_V_β_3_] ranged from 50 nM to 0.21 nM. The black lines are global fits to the data using a first-order 1:1 binding kinetic model for determining the association rate constant, *k*_*a*_, and dissociation rate constant, *k*_*d*_. (b) Thiol ratio study in which [α_V_β_3_] = 50 nM was introduced over chips prepared with thiol ratios ranging from 25:1 to 250K:1. The vertical dashed lines in (a) and (b) separates the association phase (left) in which the integrin solution is flowing over the surface from the dissociation phase (right) in which running buffer flows over the surface.

It was important to demonstrate that the SPR concurrent control approach has the sensitivity to measure ligand activity across a range of densities both necessary to trigger cell adhesion and commonly encountered in the extracellular environment. Assuming the formation of the two-component SAM on the Au surface follows a homogenous Poisson adsorption process gives a relationship in which the ligand surface density, *ρ*, is proportional to the inverse squared of the mean nearest neighbor distance, ⟨*r*⟩: *ρ*∝ ⟨*r*⟩^−2^ (Supporting Information). From this relationship, the SPO:SPC thiol ratios and their associated ⟨*r*⟩ were: 25:1 (2 nm); 250:1 (6 nm); 2500:1 (19 nm); 25K:1 (59 nm) and 250K:1 (187 nm). The middle of this range covers the most commonly cited spacing required for the formation of focal adhesions (60 nm)^6^, and we show below that the 250K:1 (187 nm) value significantly impeded cell adhesion to the same extent as the SPO-only control, indicating an upper bound. SPR chips prepared with each SPO:SPC ratio were tested for their response to the presence of 50 nM α_V_β_3_ (Fig. 2b). Even the lowest response at ≈50 RU for the 250K:1 chip is over 25-fold greater than the instrument’s limit of detection, demonstrating the technique is more than sensitive enough to characterize chips prepared with length scales appropriate for cell adhesion studies.

Figure 3 shows two possible approaches by which the response curves can be parameterized for summarizing ligand activity. One possibility is to record the response value at the end of the association phase, R_E_ (dotted line, Fig. 3a inset), as indicative of ligand activity. Another option is to measure the response slope early in the association phase, which to first order is predicted by law of mass action kinetics to be linear^35^. This was indeed the case for cRGD/α_V_β_3_ pair (Fig. 3b, inset). This second parameterization approach has the advantage of a more intuitive interpretation somewhat analogous to Michaelis-Menton enzymatic activity measurements. In the enzymatic case, the rate of product formation is plotted versus substrate concentration, while in this assay the rate of receptor-ligand pair formation is plotted against a ratio proportional to active ligand surface density (Fig. 3b, main plot). The main plots of Fig. 3a and Fig. 3b consist of at least 5 pooled parameter values for every SPO:SPC ratio, acquired using 3 independently prepared SPR chips.

**Figure 3.**
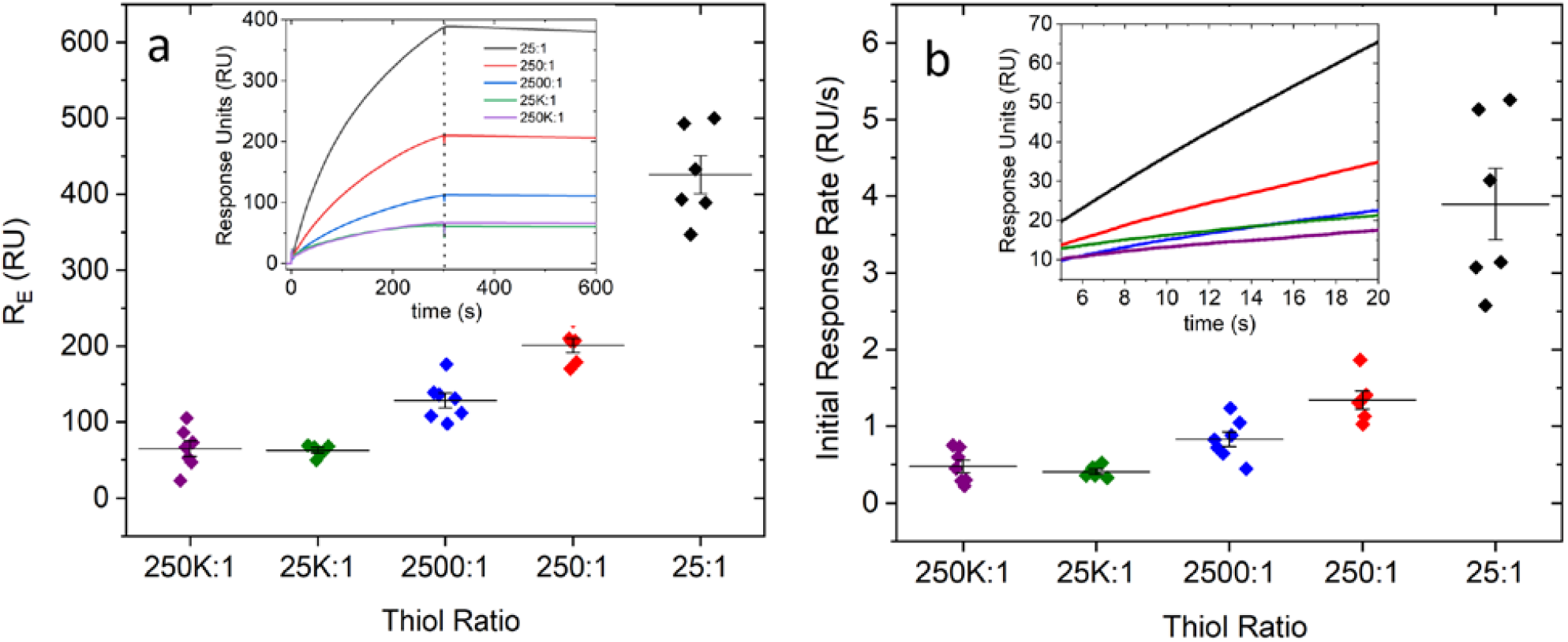
Surface ligand activity parameterization. (a) Parameterizing cRGD activity by the response unit (RU) value at the end of the association phase, R_E_. The inset is a reproduction of Fig. 2b to illustrate the demarcation of R_E_ by the vertical dashed line. (b) Parameterizing cRGD by the initial linear response rate just after injection in the association phase. The inset shows the linear response for a range of thiol ratio samples while the main figure plots the slopes derived by linear regression. The main plots of (a) and (b) consist of at least 5 pooled parameter values for every SPO:SPC ratio, acquired using 3 independently prepared SPR chips. Each color is associated with a given thiol ratio (*e*.*g*. black – 25:1). The mean and standard error bars are superimposed on the data.

The protocol described above meets the most basic requirements for a concurrent activity control. First, it produces parameters proportional to surface ligand activity using a technique (SPR) established as reproducible across multiple labs and a variety of instrument manufacturers^31^. Second, as we have previously shown, the same type of SPR studies can be used to screen for cell media factors which can significantly diminish surface activity, and determines quantitatively the extent of diminished activity^30^. This enables surface activity characterizations that include all components of the *in vitro* study but the cell itself. Finally, we have shown that the instrument has the sensitivity to characterize the range of densities most appropriate for *in vitro* cell biology. This last point would be even more useful if we could directly measure the active ligand density rather than inferring it from theoretical calculations as described above and the *in vitro* cell responses (see below). Although beyond the scope of the current study because it typically involves combining SPR results with those from additional ligand quantification techniques (e.g. radiolabeling, fluorescence), such extensions of the current technique are certainly possible and will be discussed in more detail below.

We now turn to the live cell microscopy data collected on the simultaneously prepared *in vitro* chips. Cells were added to functionalized chips with varying thiol ratios and their morphological dynamics tracked via phase contrast microscopy as a function of time. Figs. 4a and 4b show the time dependence of the average cell area and cell circularity 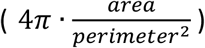, respectively. Each datum is the average of >200 cells, automatically segmented using SSML software, and pooled from the technical and biological replicates. The cRGD surface activity of the *in vitro* chips was determined by the response of the corresponding SPR control chip, enabling us to verify that the functionalization protocol was reproducible from batch to batch. Furthermore, if cellular phenotypes diverged significantly between biological replicates, while the corresponding SPR activity controls were reproducible, we were more readily able to pinpoint the cause as related to cell culture or cell processing rather than surface functionalization. The area and circularity effect sizes are maximum for the 250:1 ratio and minimum at 250K:1 (which matches the negative SPO-only control). The fact that the SPR instrument is sensitive to the recombinant integrin analyte throughout this ratio range as well is another reason the pairing of the two techniques is synergistic.

**Figure 4.**
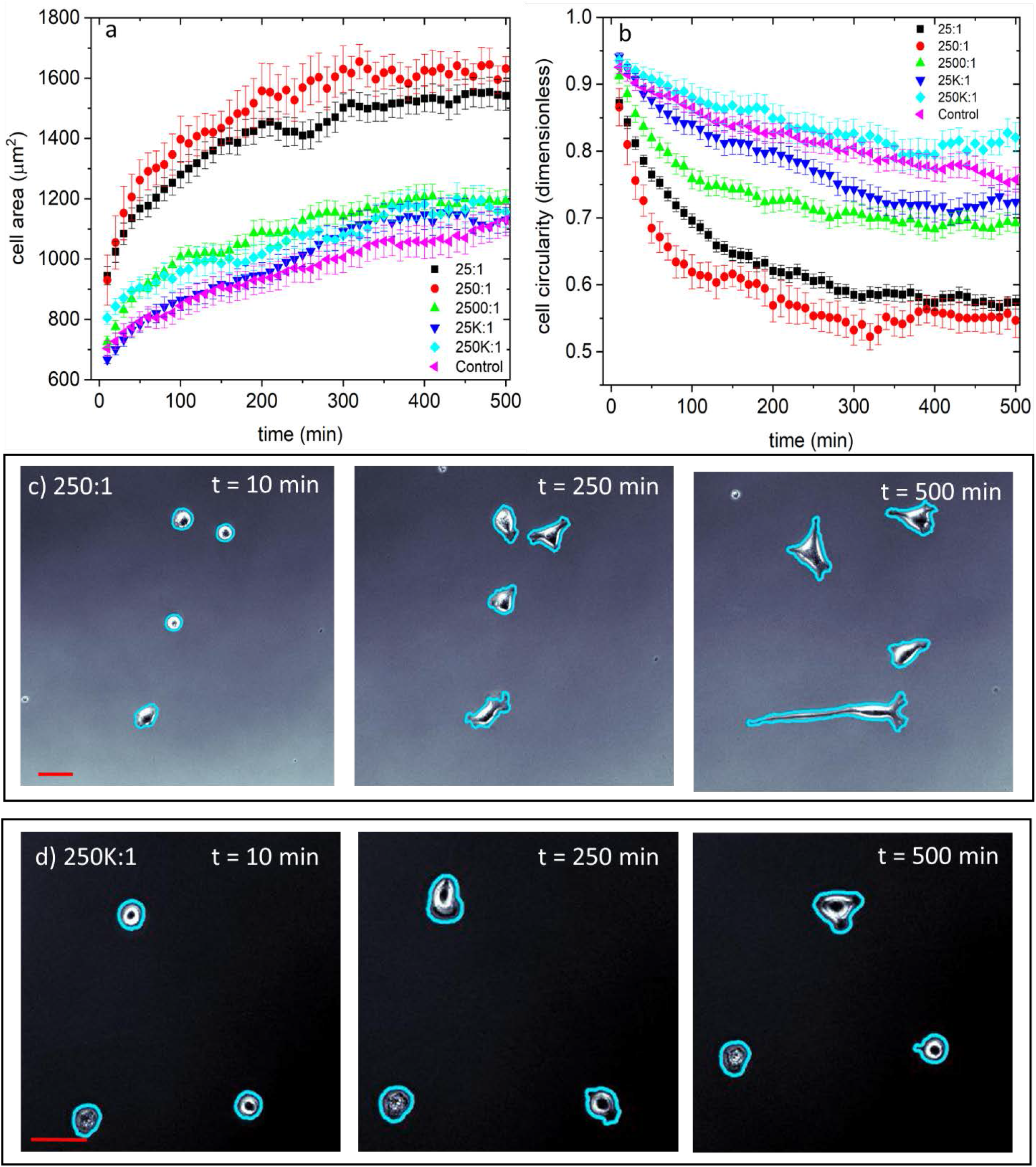
Live cell morphological responses. Cell area (a) and cell circularity (b) as a function of time for thiol ratios ranging from 25:1 to 250K:1. Representative images are shown for a 25:1 ratio chip (c) and a 250K:1 ratio chip (d) at times t = 10, 250 and 500 min. The blue outlines in (c) and (d) are the outputs of the automated SSML algorithm utilized for the area and circularity calculations. Images have been contrast enhanced to highlight cell shape. Each datum shown in (a) and (b) consists of the average and standard error from >200 segmented cells, pooled from technical and biological replicates.

The pairing of SPR and live cell microscopy can also provide insights into cellular adhesion mechanisms. Given the vast differences between the complex machinery of cellular adhesion and a SPR binding experiment, the shapes of the cell area versus time curves in Fig. 4a and the analogous SPR time course responses in Fig. 3b are remarkably similar. The cellular adhesion mechanisms, however, become more evident as the cRGD ligand density increases, with maximum cell areas achieved at the 250:1 ratio and even a reversal at 25:1. This could be due to any number of cell specific factors including focal adhesion structure, the limited number of integrin receptors available, or down regulation of integrin activity, to name a few. The SPR response conversely, which has no such constraints, continues to increase from the 250:1 to the 25:1 ratio. Figure 5 summarizes this divergence between the two types of experiments. Fig. 5a plots the initial SPR response rate from the first 15 seconds of association as a function of thiol ratio. The slope increases monotonically from the 250K:1 ratio (lowest ligand density) to the 25:1 (highest ligand density) ratios. Figs. 5b and 5c plot the initial cell area spread rate and circularity change rate from the first 100 minutes of Figs. 4a and 4b, determined by linear regression. The area and circularity slopes are also monotonic functions of the thiol ratio until 25:1, at which point there is a reversal.

**Figure 5.**
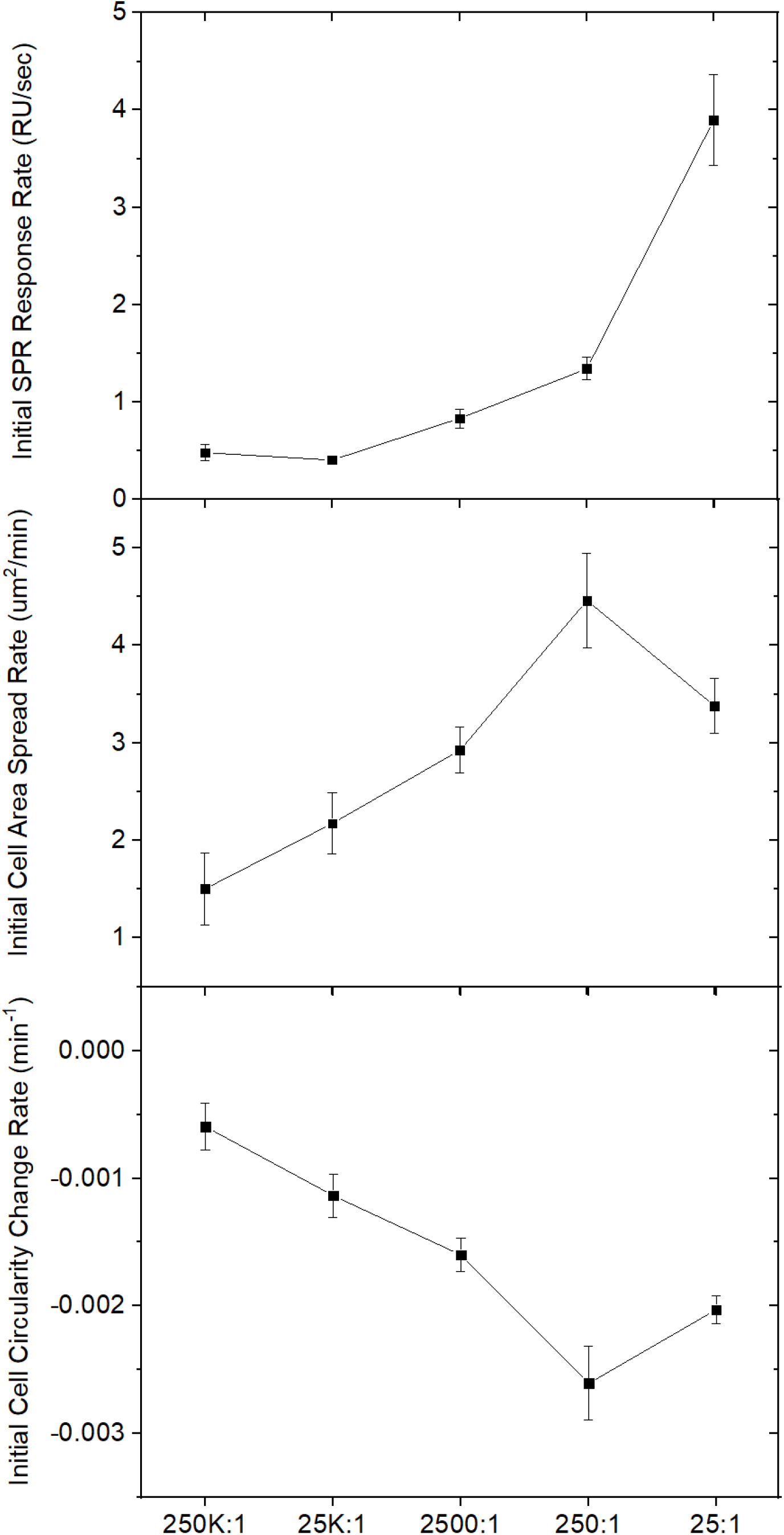
Biomolecular kinetics versus cell morphological kinetics. (a) The initial SPR response rate as a function of thiol ratio. Cell area (b) and circularity (c) response rates as a function of thiol ratio. The rate data was calculated by linear regression on the data pooled for the curves shown in Figs. 3b and 4a,b at time ranges from 5 to 20 sec and 10 to 100 min, respectively. Each datum displays the mean and standard error from the pooled regression results.

The SPR-based concurrent control technique presented here is designed to fill a surface characterization void in which ligand activity is typically assumed rather than measured. By biofunctionalizing *in vitro* and SPR chips in parallel, we have developed an approach similar to those used for surface roughness or stiffness characterization. The technique developed herein allows for quantitative ligand surface activity characterization using a measurement technique that is consistent from chip to chip, laboratory to laboratory, and has the sensitivity to detect a wide range of biologically relevant molecules. We have demonstrated that SPR meets these criteria using the most commonly employed peptide for cellular adhesion and migration studies, RGD. Furthermore, the SPR-based characterization approach can be used to screen for deleterious reductions in surface activity from components of cell media or blocking molecules (e.g. serum, BSA) which can alter cellular phenotype in unexpected ways^30^. While phase contrast microscopy was employed in this work, the surfaces are compatible with other commonly used optical modalities such as fluorescence, DIC, interference reflection microscopy, and transmitted light illumination^34^.

In future work we aim to determine a complementary approach which relates the ligand activity parameter, such as the slope in Fig. 3b, to a true mean spacing between active surface conjugated ligands rather than a thiol ratio. Such spacing information has been shown to govern adhesion, migration and provide important insights at the molecular level to the formation of focal adhesions^6,24,36^. The calculated spacing from the homogeneous Poisson adsorption model used in this work is a reasonable place to start but does not incorporate a number of important functionalization factors that may play a role - including interactions between SPO and SPC during self-assembly and cRGD conjugation efficiency - and requires experimental verification. While SPR can be used to determine the percentage of active surface ligands, it does not give an absolute density. There are, however, a number of other surface chemistry characterization techniques – fluorescence^19^, radiolabeling^36^, AFM^37^ – which, when combined with the SPR activity measurements, can be used to determine such a calibration.

It has become especially clear in recent years that reproducible *in vitro* results require careful surface characterization with regards to chemical, topographical, and mechanical inputs. Many *in vitro* studies which focus on intercellular signaling pathways are not designed in a way to carefully consider the extracellular impact on those pathways. Furthermore, studies which focus on engineering the extracellular environment typically do not quantify ligand activity. The technique presented here enables ligand activity characterization in a format consistent with investigations of both the extracellular and intercellular signaling pathways and therefore has wide applicability.

## ASSOCIATED CONTENT

### SUPPORTING INFORMATION

Cell growth curves and harvesting information; mean nearest neighbor model and calculation details

## ACKNOWLEDGMENT

M.C.R. gratefully acknowledges support from the National Research Council Research Associateship Program and the Jerome and Isabella Karle Distinguished Scholar Fellowship Program. Funding for this project was provided by the Office of Naval Research through the Naval Research Laboratory’s Basic Research Program.

