## Supplemental Information for "Interfacing Live Cells with Surfaces: A Concurrent Control Technique for Quantifying Surface Ligand Activity"

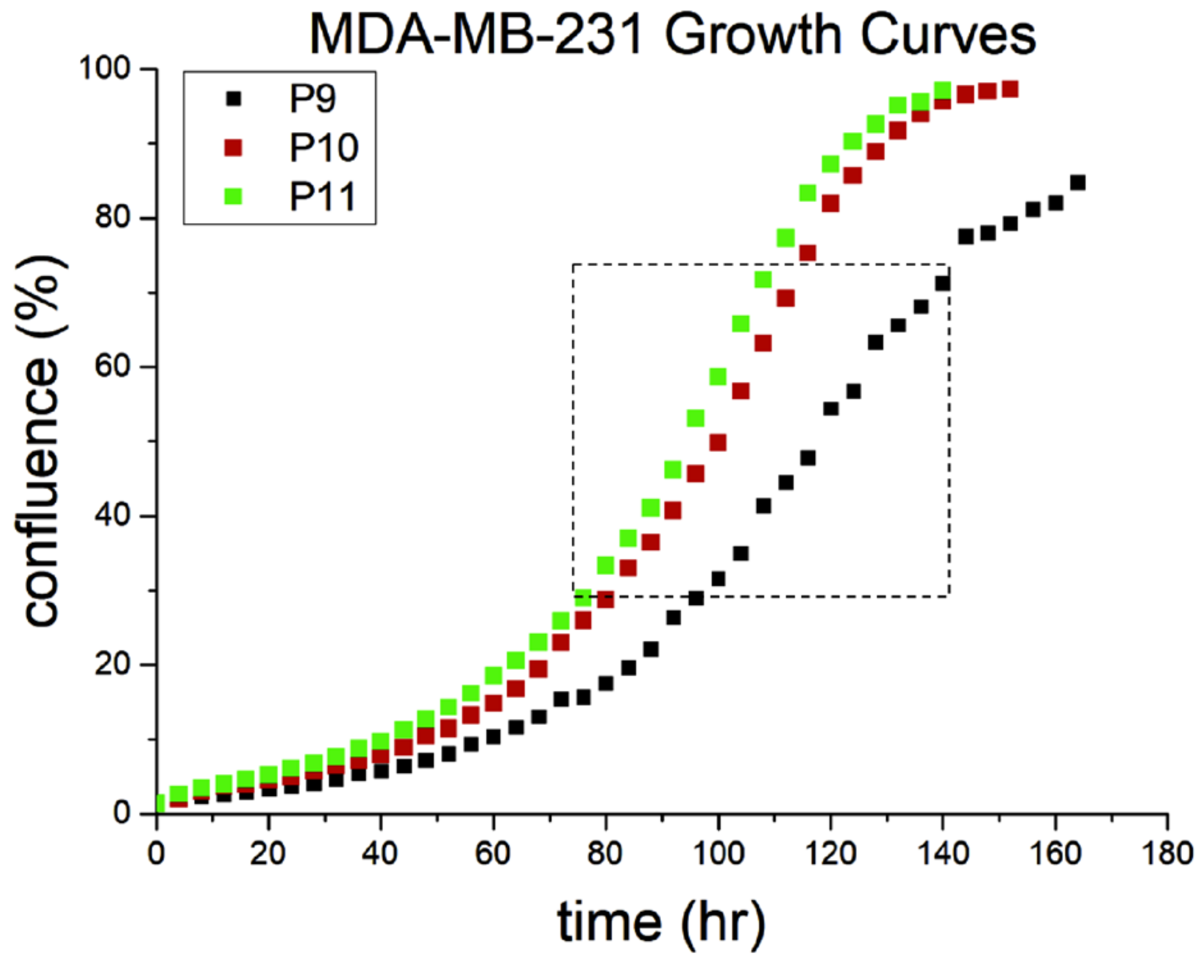


**Figure S1.** Representative growth curves of the MDA-MB-231 cells used for this study, taken via Essen Incucyte. For each experiment, a separate T25 flask seeded at similar cell density was harvested during the logarithmic growth phase (dashed box) as outlined in the main text.

**cRGD Mean Nearest Neighbor Distance Calculations**

As described in the main text, the SAM is deposited atop our gold surfaces using an ethanolic solution of SH-$(CH_2)_{11}$-$EG_6$-$COH_2$-COOH (SPC) and SH-$(CH_2)_{11}$-$EG_5$-OH (SPO) in a 1:25 ratio.

The self-assembly of thiols on Au surfaces has been extensively studied both theoretically and experimentally for decades now. Theoretically, thiol density is typically calculated based on the lattice spacing of the (111) Au surface[1]. However, given that most gold thin films used in experimental biology are polycrystalline and present a variety of orientations it is important to consider a range based on direct experimental measurements as well[2].

Given these considerations, a reasonable range of occupied area per thiolate is

$$0.22\ nm^2 < area\ per\ thiol < 0.42\ nm^2$$

The cRGD density which will be determined by the 1:25 SPC:SPO ratio, the random nature of adsorption on the gold surface during SAM formation and the conjugation efficiency. The most appropriate adsorption model is a probabilistic 2D nearest-neighbor model based on a homogeneous Poisson process, giving a probability density function of

$$g(w) = 2\rho\pi w\ exp(-\rho\pi w^2)\ ,$$

where $\rho$ is the density of SPC and $w$ is the nearest neighbor distance random variable. The mean nearest neighbor distance is then given by:

$$E[w] = 1/(2\sqrt{\rho})$$

Using the larger area per thiolate value gives the mean nearest neighbor distances quoted in the main text: 25:1 (2 nm); 250:1 (6 nm); 2500:1 (19 nm); 25K:1 (59 nm) and 250K:1 (187 nm).


1 Schreiber, F. Structure and growth of self-assembling monolayers. *Prog. Surf. Sci.* **65**, 151-256, doi:10.1016/s0079-6816(00)00024-1 (2000).

2 Nelson, K. E. *et al.* Surface characterization of mixed self-assembled monolayers designed for streptavidin immobilization. *Langmuir* **17**, 2807-2816, doi:10.1021/la001111e (2001).
